# Reporter Ion Data Analysis Reduction (R.I.D.A.R) for isobaric proteomics quantification studies

**DOI:** 10.1101/437210

**Authors:** Conor Jenkins, Alexis L. Norris, Maura O’Neill, Sudipto Das, Thorkell Andresson, Ben Orsburn

## Abstract

Isobaric labeling-based relative quantification techniques such as iTRAQ and TMT were introduced 15 years ago and are now nearly ubiquitous in shotgun proteomics labs around the world. The methods for data processing in these experiments has changed little since inception, with peptide database searching of all MS/MS spectra occurring concurrent or asynchronous to the quantification of the reporter fragment regions. In this study we present an alternative method for data processing whereby the reporter ion region of all MS/MS spectra are first examined and spectra that are not quantitatively interesting to the end user are discarded. The remaining MS/MS spectra that are retained can then be more rapidly searched for computationally expensive database alterations such as post-translational modifications and single amino acid variations in more practical time. We have termed this method Reporter Ion Data Analysis Reduction (RIDAR). To demonstrate the application of RIDAR, we reprocess a recent CPTAC 2 study containing approximately 7.8 million MS/MS spectra. Post RIDAR processing we can search this public dataset versus a human canonical FASTA database and a compiled proteogenomic database of over 875,000 known cancer mutations in a single day on a standard desktop computer, a time reduction of 85% compared to the conventional workflow. With the rapidly increasing size and density of shotgun proteomics data files, RIDAR facilitates rapid analysis of large proteomics datasets for researchers without access to high performance computational resources.

**Abstract Graphic:** 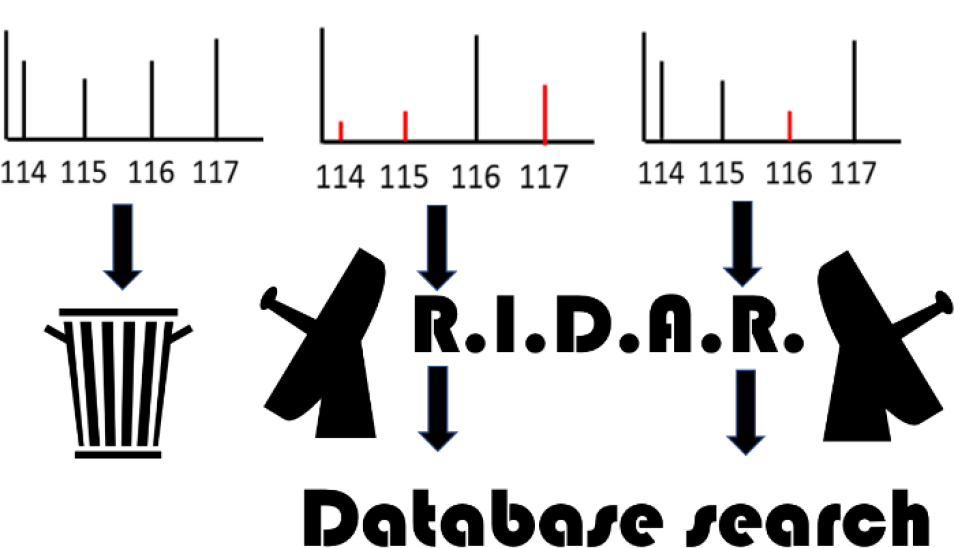

## Introduction

The analysis of shotgun proteomics data relies primarily on annotated canonical genomes that have been converted to theoretical protein FASTA sequences. These sequences are digested *in silico* to obtain monoisotopic masses and ideal fragment ion MS/MS spectra that are compared to experimental samples that are digested in the lab with the same enzymes and fragmentation methodologies. Peptides that perfectly match the canonical sequences are reasonably easy to identify with high confidence in standard proteomics workflows. Post-translational modifications and any sequence variation from the perfect *in silico* digestion, such as high numbers of missed cleavage events or single amino acid variants are more computationally difficult and often are not searched for unless the experiment absolutely requires it. While *de novo* and “delta mass” sequencing algorithms are available the massively increased computational processing time and resulting increased output size currently complicates these analyses to beyond practical for most mass spectrometry studies (1).

A common workflow for proteomic analysis involves the use of isobaric reporter ion tagging reagents such as iTRAQ and TMT (2). These reagents bind to any terminal amines, including lysines and the n-terminus of digested peptides. The mixture of labeled peptides produces virtually identical chromatographic elution patterns and MS1 signatures and are indistinguishable until fragmentation. Following fragmentation the relative quantification of the peptides in each sample is revealed by the intensity of the reporter ions in the MS/MS spectra. In a typical workflow, all MS/MS spectra are examined for peptide spectral matches (PSMs) and for quantification. Since the introduction of these technologies, there has been little change in the central strategy for data processing in these experiments. In all software that we have utilized, MS/MS spectra that pass simple quality filters are subjected to database search concurrent or asynchronous to the quantification calculations for the reporter region.

Recent advances in the scanning speed, resolution and sensitivity of shotgun proteomics instruments along with improvements in prefractionation technologies have produced a steep increase in the number of MS/MS spectra acquired per experiment and study (3). Shotgun proteomic studies containing millions of MS/MS spectra have helped identify the weaknesses in some of the classical proteomic analysis software (4) and have led to improvements in the statistics (5) and overall quality of most data analysis workflows (6).

One challenge corresponding to the relative increase in shotgun proteomic file size has been searching such large numbers of MS/MS spectra in practical time. This is particularly challenging as sample depth in some studies has shown the ability to detect post-translationally modified peptides with no secondary enrichment methodologies (7). Although a number of techniques exist for searching MS/MS data on computer clusters (8) and in cloud computing environments, it has been our observation that the majority of shotgun proteomics data appears to be processed locally on high-powered desktop computer towers. A wide gap exists in the distribution of computational power across the world, with many devices that are considered obsolete, such as 32-bit PCs, still in use in less developed countries (9). The limitations of the 32-bit architecture is well known to many scientists and was highlighted as a particular challenge for mass spectrometry as early as 2011 (10). Currently, the access to proteomics data may currently be superior to that of any other “-omics” technology thanks to organizations such as the Proteome Xchange Consortium (REF), CPTAC (REF) and NIST (https://vmsshare.nist.gov/). However, it is worth noting that the rapid increase in file size can be a significant challenge to researchers interested in pursuing protein informatics who do not have ready access to high power computational resources or the continuous access to high speed internet necessary to transfer such files (11).

In this study we describe an alternative methodology whereby the reporter ion region of all MS/MS spectra is read by a lightweight, rapid, and scalable Python script. Using user-defined tolerances and quantitative filters, the Reporter Ion Data Analysis Reduction (RIDAR) script constructs an MGF file consisting of only the MS/MS spectra that quantitatively differ according to the experimental cutoffs defined by the end user. Following RIDAR processing, we can rapidly search large datasets with multiple modifications or while using large proteogenomic databases with minimal increase in overall processing time.

## Materials and Methods

### LC-MS/MS data

All RAW files from a recent CPTAC 2 study (12) assessing 24 patient derived xenografts with iTRAQ 4-plex labeling and extensive offline fractionation were obtained from the CPTAC portal (https://cptac-data-portal.georgetown.edu/cptacPublic/). The eight 4-plex sets consist of three patient samples in each plex and a pooled control from all patients as the fourth channel. The RAW files are approximately 300GB in total size and contain approximately 7.8×10^6^ MS/MS spectra. Proteome Discoverer 2.2SP1 (Thermo Fisher) full release or freely available “viewer” version of the software was used to convert all MS/MS spectra to MGF using a vendor installed template in the software.

### RIDAR Filter

RIDAR is written in the Python language (version 3.6.3). The filter first searches for files that will be filtered by searching for .mgf extension files in the working directory. Next, user specified data residing in a standard config.txt file is used to assign the filtering parameters of RIDAR including the m/z for each reporter ion in the labeling technique, the m/z tolerance and the value of a specified fold-change between the control channel and the other channels. **Figure 1** is a flowchart demonstrating the basic RIDAR strategy.

**Figure 1.**
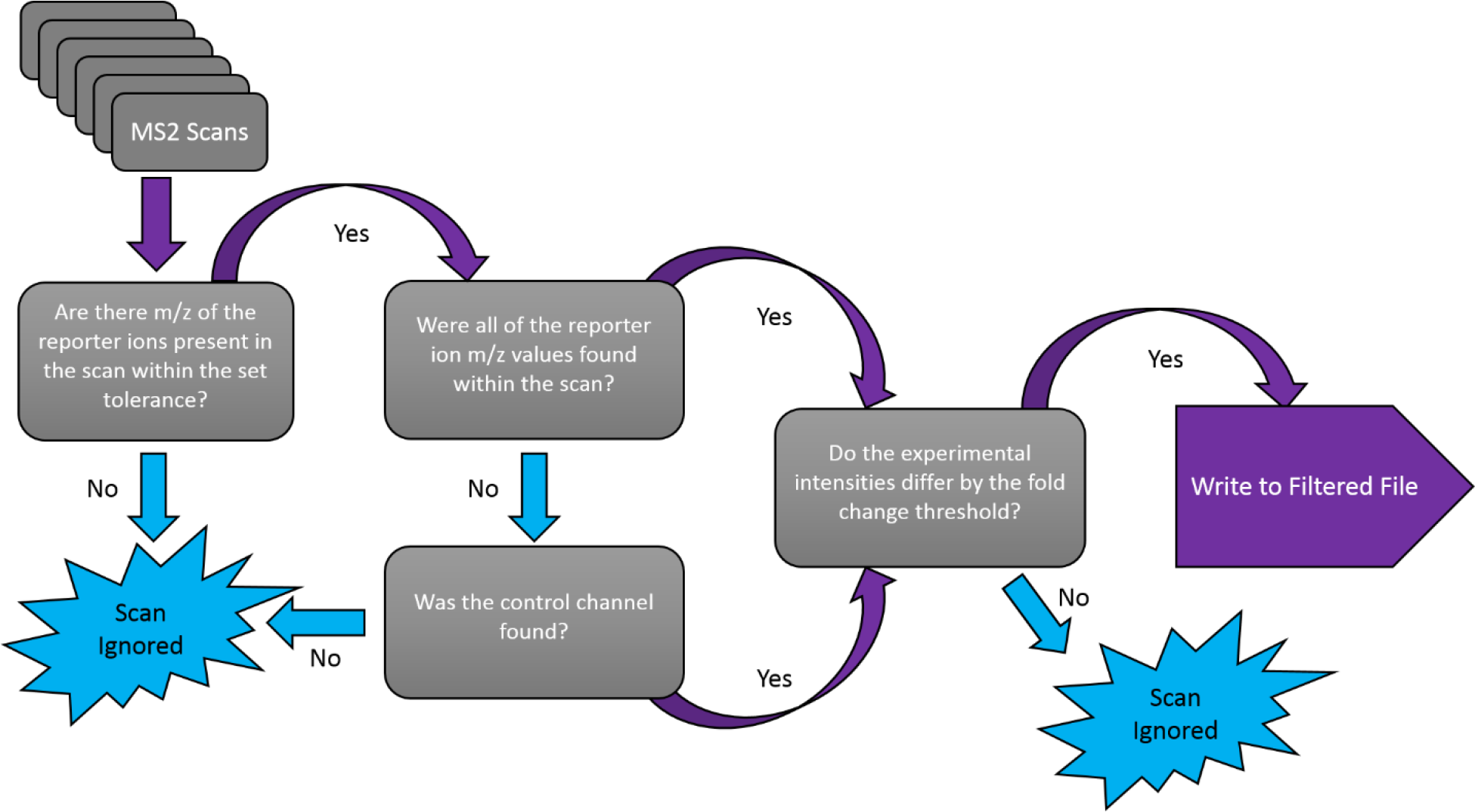
A flowchart of the RIDAR logic

RIDAR uses a matrix comparison algorithm to identify spectra of interest that fit these parameters and write them to a new filtered file without manipulating the original file. **Figure 2** describes the NxN matrix using iTRAQ 4-plex reagents as an example. An initial pass of the file of interest calculates the average intensity found for each of the specified channels across the entire MGF file. These averages are then compared to each other to populate a NxN matrix (normalization matrix figure) where N = the number of reporter ions specified in the config file. If any of the comparisons contain a value less than one, the inverse of the comparison is taken to simplify the comparison process downstream by creating a reflective, symmetric matrix. This step results in utilization of the normalization matrix in conjunction with the user specified fold-change value, producing a true fold-change threshold that the observed comparison value between channels must pass is generated.

**Figure 2.**
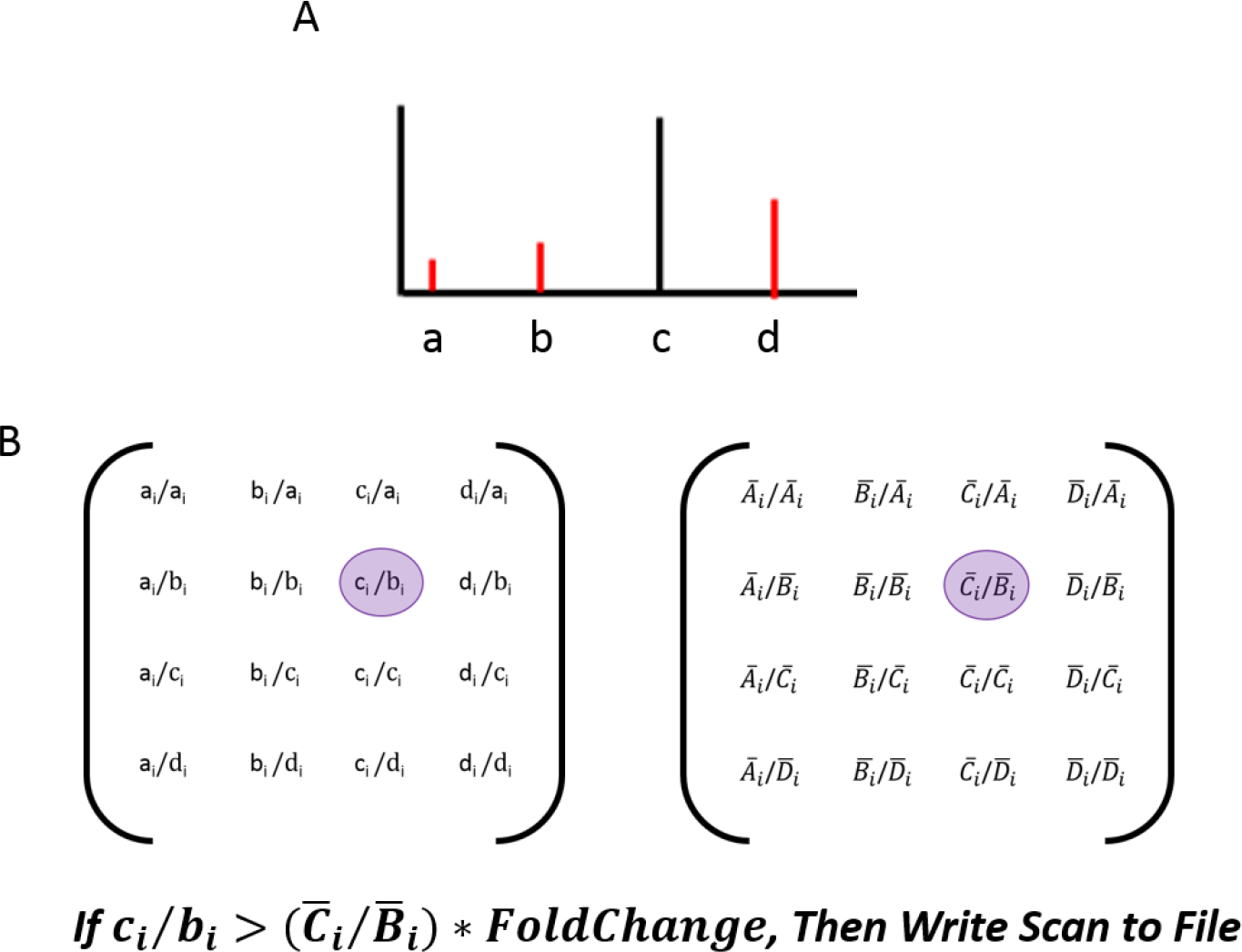
An overview of the NxN normalization process. (A) Within the scan, the reporter ions are isolated by m/z and their intensities values are collected. A matrix (the comparison matrix) is then populated with all of the possible comparisons between the channels in the same positional alignment as the associated normalization ratio between the compared channels in the normalization matrix (B). The algorithm then uses a positional comparison method to identify if any differences between the intensities of the channels exist that are greater than the specified fold change with the normalization ratio factor in the same position in the normalization matrix.

RIDAR then parses through the MGF file looking at each scan individually for the presence of reporter ions. If the control channel is present in the scan, a new matrix as shown in **Figure 2** is created containing the intensity comparison between the channels. As described above, to simplify the comparative process downstream, if the ratio is less than 1, the inverse is taken to create a reflective, symmetric matrix. If no reporter ions are in the scan within the set m/z tolerance were found, the scan does not pass the filter. If a channel is missing from the scan, a complete knockout cannot be discounted, therefore the comparisons of the intensities between that channel and the others are set a value of zero to avoid a divide by zero error.

Both matrices contain corresponding comparisons in each relative position in the matrix, as shown in **Figure 2B**. If any of the corresponding locations on the observable matrix are greater than the normalization ratio multiplied by the specified fold-change, then that scan is determined to contain aspects that pass the set parameters and will be written to the new filtered file. At the end of the filtering process, a metadata file is created detailing the total spectra in the file, the total number of spectra removed, the amount of time it took to filter and the average intensities of each of the specified channels.

The RIDAR script is open source and available at https://github.com/jenkinsc11/R.I.D.A.R.

### Computer Hardware

All database searching of the RAW, MGF and RIDAR MGF filtered data was performed on a performance desktop computer from OmicsPCs (Baltimore, MD). The system consists of an i7-6950x 10 core 3.0Ghz processor with 64GB RAM. The Windows 7 Professional operating system and data processing software are installed on a 1TB solid state drive (SSD). SSDs are relatively new storage devices that do not utilize a disk for storage, rather an integrated circuit with no moving parts. SSDs have the advantages of faster read/write speeds but are currently more expensive per unit of storage than conventional Hard Disk Drives (HDD) (13). All data processing was performed with Proteome Discoverer 2.2SP1 and data was stored during processing on either the SSD or on a 6.0TB 7200RPM HDD from Western Digital as described in the results and conclusions. All filtering of the MGF data was performed on a standard commercial Dell Latitude E7470 laptop containing an i7-6600U 2.8GHz processor with 16GB RAM and 320GB HDD using the Windows 7 enterprise operating system.

### Database searching and interpretation

The manufacturer’s default workflow for reporter ion quantification for iTRAQ 4-plex for a Q Exactive instrument was used for all analyses. This consisted of SequestHT with a 10ppm MS1 filter and a 0.02Da MS/MS tolerance. Percolator was used for PSM level FDR estimation with target decoy FDR for peptide groups and for protein group assignment. For the canonical human database, UniProt/SwissProt (downloaded February 2018) was parsed within the vendor software on “sapiens”. The cRAP database (https://www.thegpm.org/crap/) was used to flag contaminants in both the processing and consensus workflows. For mutational analysis we used the complete XMAn database (14) which consists of 872,215 translated mutated sequences compiled from COSMIC, IARC P53, OMIM and UniProt KB. The translated entries contain two missed cleavages to provide flanking peptides for each mutational region. The Proteome Discoverer “Protein Marker node” was used to flag the origination site of the FASTA match. By setting the FASTA preference to UniProt when PSMs could be attributed to both databases our final output reduced mutational flagging and annotation only occurred when sequence differed from canonical in these regions.

### Plotting and Figure Construction

Figures were generated in R (version 3.5.0) using the TidyVerse R packages (version 1.2.1; https://www.tidyverse.org/). All figure input and details on installed packages are included in the file titled “**Figure input and settings**” at the RIDAR github account listed above.

## Results and Conclusions

### File Reduction

The RIDAR filtering variables were placed in the configuration file prior to analysis. It was specified that any spectra that had at least one reporter ion mass within a 2ppm window was selected to be analyzed by the RIDAR algorithm. If an intensity of a reporter ion differed by a 2.0-fold change when compared to the control reporter ion intensity, the scan is identified as an appropriate candidate for searching, thus written to the filtered MGF file. The filtering process took less than 2 minutes per MGF file in the CPTAC study and yielded an average file size reduction of 183.88MB (~70%), correlating to an average removal of 24,854 spectra per file.

### Data processing time comparisons

Proteome Discoverer (PD 2.2SP1) divides all shotgun analysis into two sequential steps – the “Processing Workflow” for the generation of Peptide Spectral Matches (PSMs) and PSM quantification, and a second “Consensus Workflow” for compiling PSMs into peptide groups and protein/protein groups. In all iterations of data processing, searching RIDAR filtered files required a fraction of the total processing time of the original files, as shown in **Figure 3** as a sum of time for both steps. During data processing PD 2.2 creates three files. A permanent SQLite file of the PSMs, with the suffix .MSF, the permanent protein consensus output with the suffix .PDresult and a temporary intermediate file called the .Scratch. In order to process the complete 300GB of RAW data, approximately 800GB to 1.2TB of hard drive space is required to accommodate RAW files, .Scratch and final output. Due to this, it was not possible to attempt processing of the full set on the 1TB solid state C:/ drive containing operation systems and data processing software. However, the smaller RIDAR filtered files could be searched on the SSD drive and provided a further decrease in total processing time for the UniProt + XMAn experiment, decreasing from 23.0 to 15.5 total hours compared to the identical experiment performed on the HDD.

**Figure 3.**
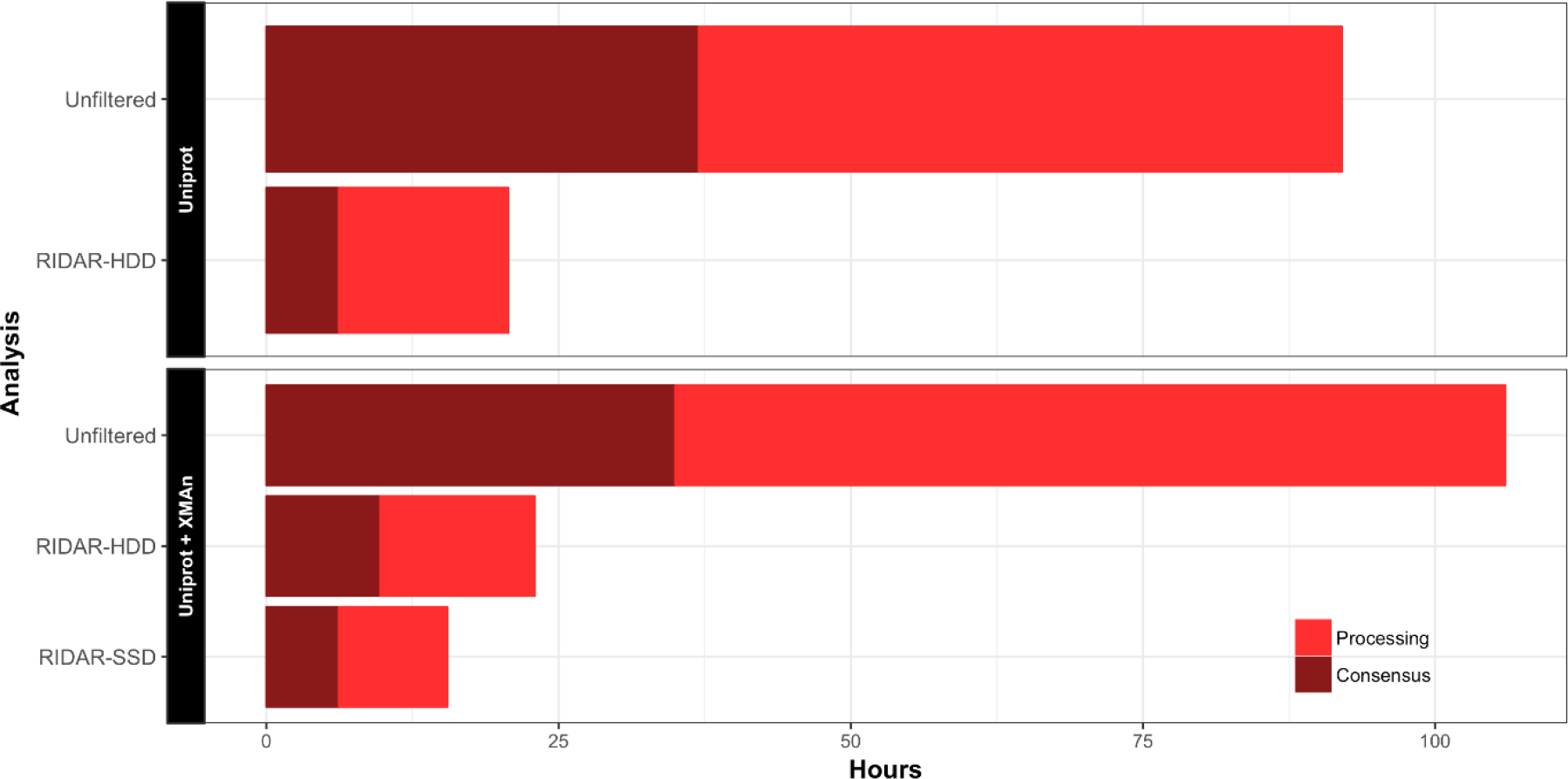
The amount of time to perform the data processing analysis in Proteome Discoverer 2.2 in hours on the original CPTAC data set (Unfiltered) and with RIDAR 2-fold filtered on Hard Disk Drive (HDD) and RIDAR 2-fold filtered on Solid Sate Drive (SSD)

### Results summary

Using the RIDAR script we can filter spectra containing reporter ions to any level that the user chooses. The file size is reduced by removing the spectra the end user finds least interesting first. When starting with this dataset, filtering to a 2.0-fold cutoff results in a reduction of 7.8×10^6^ spectra to 3.2×10^6^ spectra. Further experiments with filtering are shown in **Table 1**. Moving from a 2.0-fold RIDAR cutoff to a 5.0-fold one further reduces the number of spectra leaving less than 10% of the original starting number.

**Table 1.**
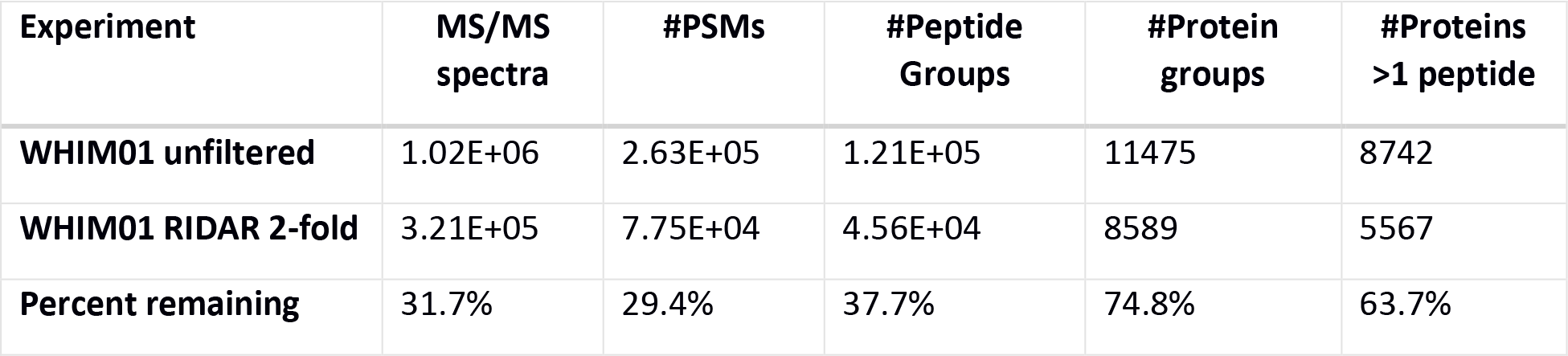
The effect of the reduction of spectra on the WHIM01 sample set

| Experiment | MS/MS spectra | #PSMs | #Peptide Groups | #Protein groups | #Proteins >1 peptide |
| --- | --- | --- | --- | --- | --- |
| WHIM01 unfiltered | 1.02E+06 | 2.63E+05 | 1.21E+05 | 11475 | 8742 |
| WHIM01 RIDAR 2-fold | 3.21E+05 | 7.75E+04 | 4.56E+04 | 8589 | 5567 |
| Percent remaining | 31.7% | 29.4% | 37.7% | 74.8% | 63.7% |

In order to determine how the RIDAR reduction affects the final results, the CPTAC file subset WHIM01 were arbitrarily selected for in-depth analysis. The four iTRAQ labels were applied to three patients and to the 117 labeled pooled control that was used for all samples. We processed these intact files against the UniProt Human SwissProt FASTA as well as the 2.0-fold RIDAR-filtered files for this set. **Table 1** demonstrates the reduction in MS/MS spectra, PSMs and peptides/proteins.

The reduction in the number of MS/MS spectra has little effect on the identification of the proteins in the smaller dataset, with 99.8% of protein groups found in the RIDAR 2.0-fold set also found in the larger unfiltered output when two or more unique peptides are used as a threshold cutoff, as shown in **Figure 3A.** The 12 proteins that appeared only in the RIDAR-filtered dataset are included in **Supplemental Table 1**.

**Figure 3A.**
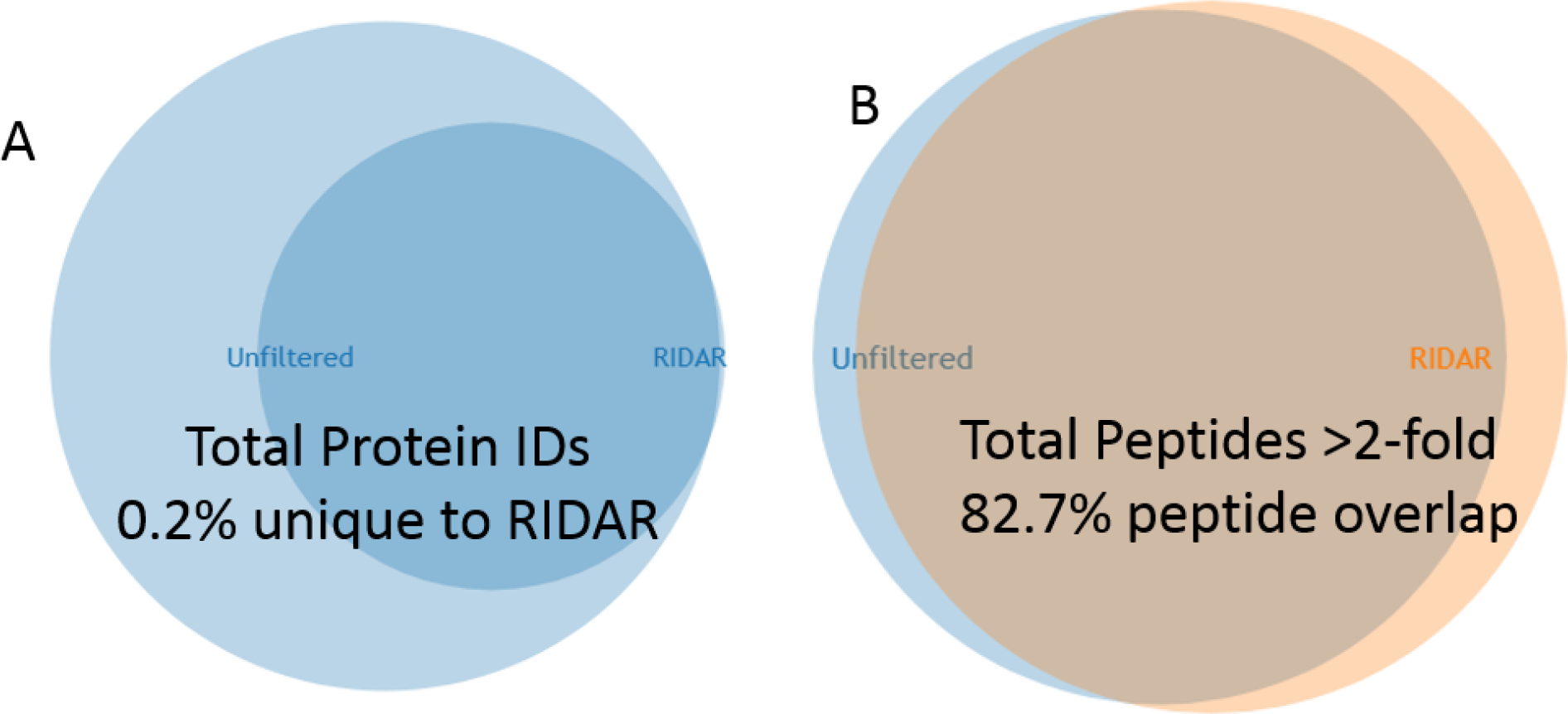
Venn Diagram of the protein group identifications in the WHIM01 vs WHIM01 RIDAR 2-fold filtered dataset, demonstrating that reducing spectra does not cause protein identifications to markedly change. **B)** A diagram demonstrating the proteins that have a ratio of 117/114 greater than 2-fold above the arithmetic mean in the two sets.

The reduction in MS/MS spectra and PSMs appears to scale with the reduction of the full set and can be readily explained by the loss of the peptides that appear to match in overall abundance across all samples. A complete overview of the result statistics of these two files are available in **Supplemental Table 2**. The decrease in percent protein coverage is the metric most affected, with a 62% drop in the number of PSMs and 48% drop in peptides per protein contributing to a 41% drop in protein coverage for each protein identified. However, the RIDAR filtered set still contains an average of 10.42 PSMs per identified protein. As it is still common practice in many proteomics data processing pipelines to quantify, at most, the three most abundant peptides in each measured protein it is possible that the loss of peptides may have little to no affect on the quantification of proteins in some analyses.

To assess potential changes in protein quantification, the protein accession numbers and quantification ratios were extracted for 114 compared to the 117 control channel from both the intact WHIM01 and RIDAR-filtered files. As expected when removing large number of quantitatively similar measurements, such as the 1:1, the average mean and standard deviation of the two sets were altered. In the unfiltered set the average ratio of 117/114 is 1.14 ± 0.76. Post RIDAR 2.0-fold filtering, this increases to 1.90 ± 1.35. To properly determine the significance of quantification in proteomics experiments, advanced statistical models such as the empirical Bayes model are the only true recourse. On top of the inherent issues in peptide relative quantification, the matching of peptide spectral matches to proteins and protein groups adds another layer of complexity to the challenge (15). To circumvent this, we compared the number of quantified peptide groups with a greater than 2.0-fold change between 117/114 in the WHIM01 full and RIDAR 2.0-fold filtered sets. Of the 9,110 peptides compared at this cutoff, 82.7% of the peptides were conserved, as shown in **Figure 3B**. For the peptides identified by only one of the two methods, 318 peptides were uniquely found within the RIDAR-filtered set, and 546 peptides were unique to the unfiltered dataset. While a perfect overlap might strengthen the case for the usefulness of the RIDAR dataset within conventional workflows, the cumulative effects of peptide identification level false discovery rates and the relative nature of proteomics quantification suggests to use that this tool still has large potential value.

## Conclusions

We have demonstrated the application of a seemingly novel pre-processing methodology for reporter ion quantification experiments. By filtering out MS/MS spectra that do not meet the end user’s criteria for significant prior to analysis, we can more rapidly process large datasets and search for mutations using proteogenomic databases. Further development on RIDAR will continue on three main fronts. The first is to allow the filtering of data derived from MS3-based proteomics workflow. The second is to make the utilization of the script simpler for the end user. Finally, we are working on the development of a cloud server that will allow a user to reduce files from public repositories automatically, enabling analysis of large datasets in institutions around the world regardless of access to high performance computational resources.

